# Individual differences in the pharmacokinetic profiling of Δ9-THC may be associated with differential motivational effects of Δ9-THC on cognitive performance

**DOI:** 10.1101/2025.09.30.679373

**Authors:** Teneisha M. Myers, Noah C. Neverette, Aaron S. Devanathan, Mary M. Torregrossa

## Abstract

**Rationale:** Cannabis is the most widely used illicit substance in the United States, and its use is increasing with the recent push for legalization and decriminalization, as well as the growing use for medicinal purposes. While cannabis can have positive effects in some individuals, there are potential negative consequences including dependence, psychosis, and cognitive impairments. The top reported reasoning for using cannabis is to alleviate stress, however, whether cannabis differentially induces positive or negative effects under stress has not been studied.

**Objectives:** The current study investigates whether stress affects working memory, and if THC can exacerbate or ameliorate its effect. An additional aim is to determine if plasma THC concentrations are associated with cognitive performance

**Methods:** Adult male and female Sprague Dawley rats performed a delay-match-to-sample working memory task. Rats assigned to the stress group were exposed to acute restraint stress prior to administration of either vehicle, 0.5,1, or 3 mg/kg THC. Blood samples were collected 5, 25, 60, and 120 minutes after administration.

**Results:** Acute restraint stress and THC did not impact working memory. However, acute administration of 3mg/kg THC disrupted motivation-related engagement in the task in a subset of rats. These non-responders exhibited greater plasma THC and metabolite concentrations compared to rats who maintained baseline response rates after 3mg/kg THC administration.

**Conclusions:** Individual differences in the pharmacokinetic/metabolic profile of THC may be associated with differential sensitivity to cognitive/motivational effects of THC and highlight one potential mechanism for the diversity of reported adverse versus positive outcomes after THC exposure.

## 1. Introduction

Cannabis and cannabis derivatives are the most widely used illicit substances in the United States^1,2^. The prevalence of cannabis use has been on the rise alongside the push for the legalization of its recreational and medicinal use. From 1992 to 2022, the number of individuals reporting consistent, past-month use of cannabis increased drastically from 0.9 million to 17.7 million^3^. Alongside this increase in prevalence, Δ-9-tetrahydrocannabinol (THC) content, the main psychoactive derivative of cannabis responsible for its rewarding effects, in today’s cannabis products is around 10 times higher than it has been historically. These recent changes in the legalization of cannabis and the emergence of higher THC content strains prompt the need for more research on the biological consequences of THC use.

In particular, the increase in legalization of cannabis altered the public’s perception of the safety of cannabis use^4–6^. Recent studies have reported that adolescents and young adults perceive cannabis as less harmful than previous generations^4,5,7^. Despite this, many individuals who use cannabis experience adverse consequences; approximately one in three individuals using cannabis develop issues related to its use that impair normal functioning^8^. Negative consequences following cannabis use can include the development of a cannabis use disorder (CUD); with more people seeking treatment for cannabis use disorder than for any other illicit substance, psychotomimetic effects in genetically predisposed individuals, as well as impairments in neurocognitive functioning^2,8–11^. Although negative effects following cannabis use have been reported, it is also clear that for others, cannabis use is beneficial. Previous studies have demonstrated its clinical benefits for the treatment of anxiety, chronic pain, insomnia, and chemotherapy-induced nausea^12^. With the ability of cannabis to produce both positive and negative effects following its use, the need to develop a deeper understanding on factors that give rise to these individual differences is necessary.

Factors that are widely known to influence individual differences in the effects of a drug are their pharmacokinetics. The pharmacokinetic profiling of a drug plays a critical role in understanding drug behavior through its absorption, distribution, metabolism, and excretion^13^. In rodents, sex-based differences in the pharmacokinetics of THC have been previously reported^14^. In humans, sex differences are also apparent, with males showing greater peak plasma THC concentrations following inhalation of high-potency THC, while females exhibit higher plasma THC concentrations after oral administration^15^.

While previous studies have investigated the pharmacokinetics of THC, there is a gap in the literature mapping pharmacokinetic profiling on to behavioral endpoints^16^. Another factor potentially mediating individual differences in the response to cannabis use that has been overlooked in the field is the physiological state of the individual at the time of consumption. To date, the most commonly reported reason for the initiation of cannabis use, as well as its continued maintenance, is to alleviate feelings of stress^17,18^. Indeed, acute cannabis use can decrease feelings of stress and anxiety ^17,19,20^. Moreover, being in a state of stress has been shown to modulate the pharmacokinetics of drugs of abuse^21^. Stress exposure can lead to changes in blood flow rate as well as vascular function which leads to alterations in the distribution and pharmacokinetic profiling of the drug ^21^. Yet, this has not been taken into consideration when exploring individual differences in the effects of THC. Therefore, it is unknown whether the differential sensitivity to the impairing effects of THC is associated with differences in the pharmacokinetics of THC or by states of stress. A deeper understanding of individual pharmacokinetic responses will allow us to gain a better idea of how cannabis can uniquely impact different populations, as individual variations in cognitive effects of cannabis have been overlooked ^22^. The current study sought to explore whether stress differentially affects the pharmacokinetic profile of THC and if there is a relation to cognitive functioning. We hypothesized that acute restraint stress would lead to impairments in working memory performance, and that this behavioral outcome would map to changes in the pharmacokinetic profiling and metabolism of THC.

## 2. Methods

### 2.1 Subjects

Adult male (n = 24) and female (n = 24) Sprague Dawley rats (postnatal day 70 upon arrival) were purchased from Envigo (Indianapolis, IN). Animals were group-housed upon arrival and throughout the experiment in a temperature and climate-controlled facility on a 12:12-hour light/-dark cycle, with all experiments conducted during the light cycle. Rats were habituated to the animal facility for at least 7 days before the experimental training began. 24 hours before training began, rats were food-deprived. Throughout the experiment, rats were allowed access to water ad libitum and were fed 12-14 g rat chow per day immediately after the working memory task was completed. All procedures were approved by the University of Pittsburgh Institutional Animal Care and Use Committee and followed the National Institutes of Health Guide for the Care and Use of Laboratory Animals.

#### Apparatus

Behavioral training and testing were conducted in standard operant conditioning chambers housed within sound-attenuating cabinets (Med Associates, St. Albans, VT, USA). The chambers were configured with a panel containing 5 nose poke apertures on 1 wall of the chamber, a house light, a sucrose pellet dispenser, and a magazine on the opposite wall.

### 2.2 Drugs

Δ-9-Tetrahydrocannabinol (THC) was provided by the National Institute on Drug Abuse’s Drug Supply program. Stock solutions were prepared by adding 100–200 μl of Tween 80 to an aliquot of THC solution before the ethanol was evaporated off using a steady stream of nitrogen gas. The stock solution (50mg/mL) was then brought to a volume of 1 mL with sterile 0.9% saline. Working solutions were prepared immediately prior to each self-administration session by diluting the stock solution with additional saline to reach the desired concentration of THC. Vehicle solutions were made from Tween 80 with an equivalent volume dilution in saline.

### 2.3 Working Memory Training

All rats were trained to perform a delay-match-to-sample working memory task over 6 increasingly difficult stages. In stage 1, rats were trained to respond to any of the 5 illuminated apertures to receive a sucrose pellet reward on a fixed-ratio 1 (FR1) schedule of reinforcement. In stage 2, rats learned to respond into a specific illuminated aperture to receive the sucrose pellet reinforcer. In stage 3, rats learned to respond into a single illuminated aperture during the “sample” phase. After a 0.5-second delay, the “choice” phase began where the rat had to respond into the originally illuminated aperture from a choice of 3 adjacent illuminated apertures to receive the sucrose pellet reward. Once rats met criteria of 80% correct responses during this stage, the delay between the sample and choice phases was increased over the final 3 stages. Rats experienced 7 delays ranging from 0.5 to 6 seconds in stage 4, 0.5 to 12 seconds in stage 5, and 0.5 to 24 seconds in stage 6. Each delay was presented randomly within a block of trials, and all delays were presented before a new trial block began. Rats remained in stages 2–6 for a minimum of 3 days and performed at least 80% correct trials at the 0.5-second delay before advancing.

### 2.4 Drug Treatment

Immediately following completion of initial working memory behavioral training, using a latin-square design, rats received intraperitoneal (i.p) injections of THC (0.5, 1, or 3 mg/kg) or vehicle on four working memory test days (Figure 1A). Administration of THC or vehicle occurred 30 minutes prior to the working memory testing. Rats in the stressed group received THC or vehicle immediately following stress exposure.

**Figure 1.**
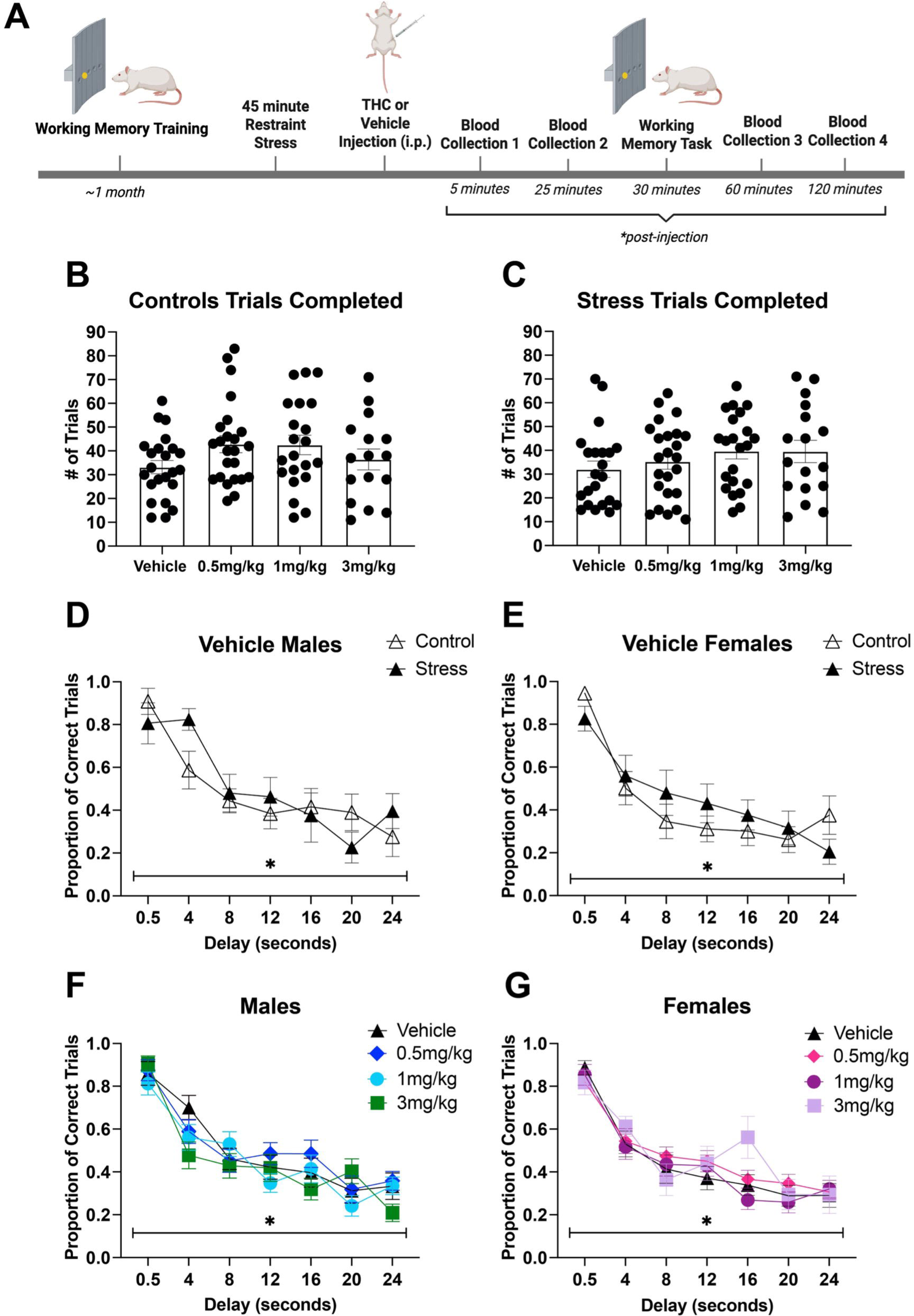
The effect of acute restraint stress and acute THC administration on performance accuracy on the working memory task. A) A schematic of the experimental timeline. Following working memory training, rats were exposed to 45 minutes of restraint stress (or home cage controls) immediately followed by THC or vehicle administration, with working memory testing 30 minutes post-injection. Blood collections were conducted at 4 timepoints following THC administration for later THC plasma pharmacokinetic analysis. B) Number of trials completed on the working memory task at each dose in the non-stressed control rats and, C) stressed rats. D) Stress does not impact working memory performance in males or E) females. F) Acute administration of THC at all doses and vehicle does not impact working memory performance in males, or G) females. All figures are mean (±SEM), main effect of delay ** *p* < .001

### 2.5 Acute Restraint Stress

To explore whether acute stress led to an alteration in working memory performance, male and female rats in the stress group were exposed to 45 minutes of restraint stress 30 minutes prior to the working memory task. Rats were restrained using a plastic restraint cone, DecapiCone (Braintree ScientificDC200).

### 2.6 Quantification of THC, 11-OH-THC, and 11-COOH-THC

Blood samples were taken via tail nick from a subset of rats (12 males, 12 females) at 5, 25, 60, and 120 minutes after THC was administered. THC, 11-OH-THC, and 11-COOH-THC concentrations were quantified in plasma using ultra-performance liquid chromatography-tandem mass spectrometry (UPLC-MS/MS). Briefly, a UPLC-MS/MS system consisting of a Thermo Scientific Vanquish UPLC and Thermo Scientific TSQ Altis equipped with a heated electrospray ionization source was used. The parent to product ion transitions used for quantitation were *m/z* 315.25 →193.05 for delta-9-THC, *m/z* 318.40 →196.1 for delta-9-THC-d3, *m/z* 331.23 →193.05 for 11-hydroxy-delta-9-THC, *m/z* 334.23 →196.05 for 11-hydroxy-delta-9-THC-d3, *m/z* 345.2 →299.1 for 11-nor-9-carboxy-delta-9-THC and *m/z* 348.2 →196.05 for 11-nor-9-carboxy-delta-9-THC-d3. Chromatographic separation of the samples was accomplished with a Waters Acquity BEH C18 (2.1 ′ 100 mm; 1.7 mm particle size) along with the BEH C18 VanGuard precolumn (2.1 x 5 mm) with an elution using water with 0.1% formic acid (A) and acetonitrile with 0.1% formic acid (B) at a flow rate of 300 µL/min. The linear range of the standard curves for each analyte was 0.1-100 ng/mL (r^2^ > 0.9986) with interassay precision and accuracy deviations <15%.

### 2.7 Statistical Analysis

Behavioral data was analyzed using GraphPad Prism version 10. Proportion of correct trials, plasma THC and metabolite concentrations in responders vs non-responders (described below) were analyzed by 2- or 3-way repeated measures ANOVA (α=0.05) followed by Bonferroni post-hoc comparisons when appropriate. Number of trials completed were analyzed using a one-way ANOVA followed by Bonferroni post-hoc comparisons when appropriate. Pharmacokinetic analysis for THC and its metabolites was performed in Phoenix 64 version 8.3.5.340 (Certara, Inc. Radnor, PA). Area under the curve (AUC) estimates were obtained through non-compartmental analysis using linear up-log down method. Data visualization, statistical analyses, and retrieval of descriptive statistics were performed in R version 4.3.3 (R Foundation for Statistical Computing (Vienna, Austria)). THC, 11-OH-THC, and 11-COOH-THC concentrations below the lower limit of quantification (BLQ) were treated as missing in non-compartmental analyses and were visually represented as half the lower limit of quantification (e.g. 0.05 ng/mL) in figures. Differences in AUC estimates between binary groups (male versus female; stress versus no stress) were determined via Wilcoxon rank-sum tests. Differences in AUC estimates – stratified by sex and stress – were determined by the Kruskal-Wallis test followed by Dunnett’s test. P-values were considered statistically significant at α=0.05, which was adjusted using the Bonferroni correction procedure for multiple comparisons.

## 3. Results

### 3.1 Acute restraint stress does not impact working memory performance

A two-way ANOVA revealed that in both males and females, acute exposure to restraint stress did not impact working memory performance, as non-stressed and stressed rats exhibited similar behavior. In both groups, at the short delay, all rats responded with around 80% or greater accuracy. From here, as the delays between the sample and choice phases became longer, the performance accuracy decreased to almost chance levels, with the proportion of correct trials completed decreasing in the males [effect of delay: F(6,66) =14.46, *p*<0.0001] and females [effect of delay: F (6,66) = 16.48, *p* < 0.0001]. There was no effect of group [F (1,11) = 0.1595, *p* = 0.6973] or a delay by group interaction [F (6,45) = 1.622, *p* = 0.1631; Figure 1D] present in the males or the females [effect of group: F (1, 11) = 0.3138, *p* = 0.5866; delay by group interaction: F (6,66) = 1.268; *p* = 0.2843; Figure 1E].

Pharmacokinetic profiling analysis revealed that plasma concentrations were similar between stressed and non-stressed groups (Figure 4B,D,F). There were no statistically significant differences in AUC estimates between stressed and non-stressed groups for THC or either metabolite in all THC dose groups tested (Figure 4B,D,F; all p-values > 0.05). After stratifying by both sex and stress, significant differences in AUC estimates for THC in the 1 mg/kg dosing group and for 11-COOH-THC in the 3 mg/kg dosing group were found. AUC estimates for stressed females were significantly higher than those observed in stressed males for THC in the 1 mg/kg dosing group and 11-COOH-THC in the 3 mg/kg dosing group (Supplemental Figures 4A,C). Concentration vs. time curves for THC and both metabolites in each dosing group are shown in Supplemental Figures 2,3.

### 3.2 Acute THC administration does not alter working memory performance

Due to the lack of stress effect observed, non-stressed controls and stressed rats were combined for the remainder of behavioral analysis. We then examined whether acute THC administration at a low, moderate, or high doses impacted cognitive performance on the working memory task. 2-way ANOVA revealed that THC did not alter working memory performance at any of the doses administered compared to the vehicle treatment. This lack of effect was present in both the males [effect of dose: F (3,69) = 1.188, *p* = 0.3207; Figure 1F] and females [effect of dose: F(3,69) = 0.7333, *p*= 0.5357]. Acute administration of THC and vehicle led to a decrease in proportion of correct trials completed as the delays became longer in the males [effect of delay: F (6, 138) = 48.06, *p* < 0.0001; Figure 1F] and females [effect of delay: F (6,138)= 42.40, *p* < 0.0001; Figure 1G]. There were no dose by delay interactions in males [F (18,330) = 1.403; *p* = 0.1273] or females [dose by delay interaction: F (18,337) = 1.075, *p* = 0.3758]. The overall number of trials completed on the working memory task was also examined, and no differences in the total number of trials completed was observed in the controls [F (3, 80) = 1.776, *p* = 0.1585; Figure 1B] and stressed rats [F (3, 81) = 1.023, *p* = 0.3869; Figure 1C] at each dose of THC administered. Rats were excluded from behavioral analysis if they did not meet inclusionary criteria of 11 or more trials completed on the test day.

THC AUC estimates were statistically significantly higher in females compared to males in the 1 mg/kg dosing group (Figure 4A). AUC estimates for 11-OH-THC and 11-COOH-THC were both significantly higher in females compared to males in the 3 mg/kg dosing group (Figure 4C,E). AUC estimates for 11-OH-THC were also significantly higher in females compared to males in the 1 mg/kg dosing group (Figure 4E). Dose proportional increases in AUC estimates were observed from 0.5 mg/kg to 3 mg/kg for THC, 11-OH-THC, and 11-COOH-THC. Concentration vs. time curves for THC and both metabolites in each dosing group are shown in Supplemental Figure 1.

### 3.3 3mg/kg THC alters responsivity on the working memory task

Following initial behavioral analysis, we discovered that there was a significant subset of rats that did not meet inclusionary criteria of at least 10 trials completed after administration of THC. Thus, we were interested in factors that led some rats to meet inclusion criteria vs rats that did not. To do so, the number of trials completed by all rats were analyzed at each dose, and a one-way ANOVA showed that acute THC administration significantly impacted the number of trials completed [F (3, 188) = 4.169, *p* = 0.0069]. Bonferroni’s post hoc test revealed that rats exhibited a significant decrease in trials completed on the working memory task following acute 3mg/kg THC administration compared to the 0.5mg/kg [t (188) = 3.13, *p* = 0.0121]. There was also a trend for a significant decrease in trials completed following 3mg/kg administration relative to 1mg/kg [t (188) = 2.65, *p* = 0.0528] (Figure 2A). From here, the proportion of rats meeting inclusion criteria (responders) and rats that did not meet criteria (non-responders) at each dose of THC were analyzed by stress group (Figure 2B). Analysis revealed that acute administration of 3mg/kg THC led to a greater proportion of non-responders in both control and stressed-exposed rats.

**Figure 2.**
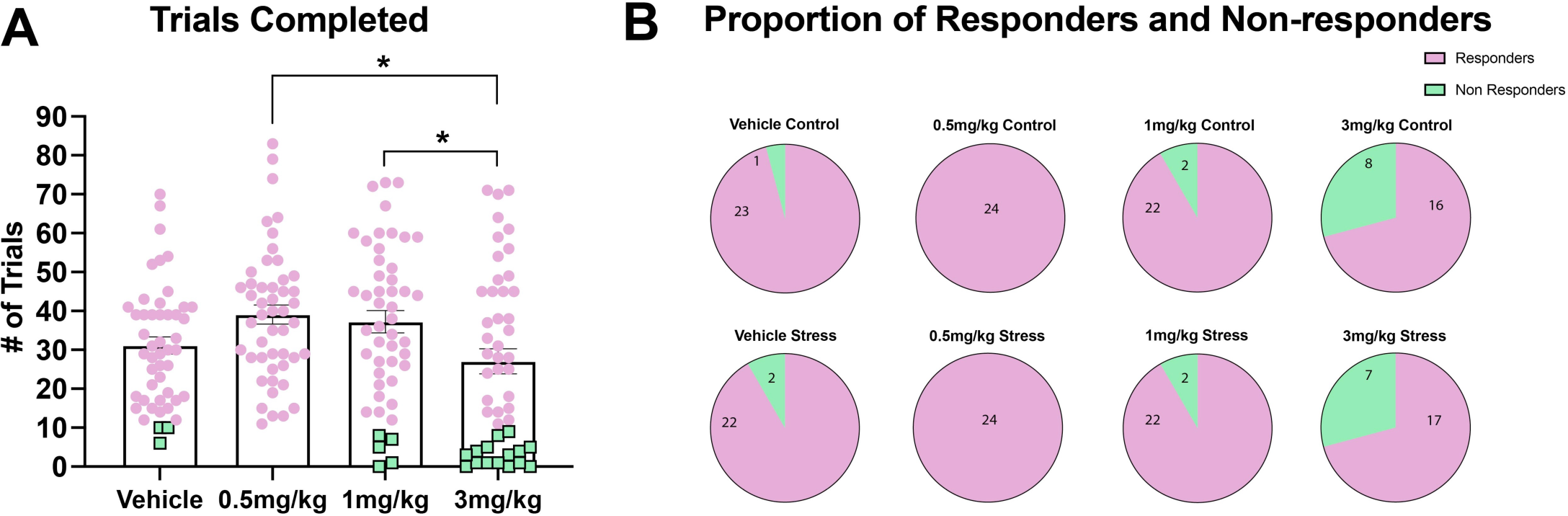
Distribution of responders and non-responders on the working memory task following THC administration. A) Number of trials completed by responders (purple circles) and non-responders (green squares) following each dose of THC, data represented as mean (±SEM), * *p* < .05, B) Proportion of responders and non-responders by stress group following each dose of THC.

### 3.4 Non-responders exhibit elevated plasma THC and metabolite concentrations following acute 3mg/kg THC administration

We next asked whether there was a relationship between responsivity on the working memory task and plasma THC and major metabolite concentrations following 3mg/kg THC administration. Previous research has shown that there are sex differences in THC plasma concentrations following i.p injection^14,15,23,24^. Based off this, we first determined the pharmacokinetic and metabolic profiles of THC and its major metabolites of the responders and non-responders stratified by sex (Figure 3). A 3-way ANOVA revealed a main effect of time [F (3, 51) = 4.027, *p =* 0.0120] and a time by responders vs non-responders interaction [F (3, 51) = 5.798, *p* = 0.0017] in the THC plasma concentrations, with no effect of sex (Figure 3A). Time course plasma concentrations of 11-OH-THC revealed a main effect of responders vs non-responders [F (1, 19) = 5.999, *p* = 0.0242] and a main effect of males vs females [F (1, 19) = 15.44, *p* = 0.0009; Figure 3C]. Analysis of 11-COOH-THC plasma concentrations revealed a main effect of time [F (3, 48) = 6.990, *p* = 0.0005] and a time by males vs females interaction [F (3, 48) = 4.278, *p* = 0.0094; Figure 3D].

**Figure 3.**
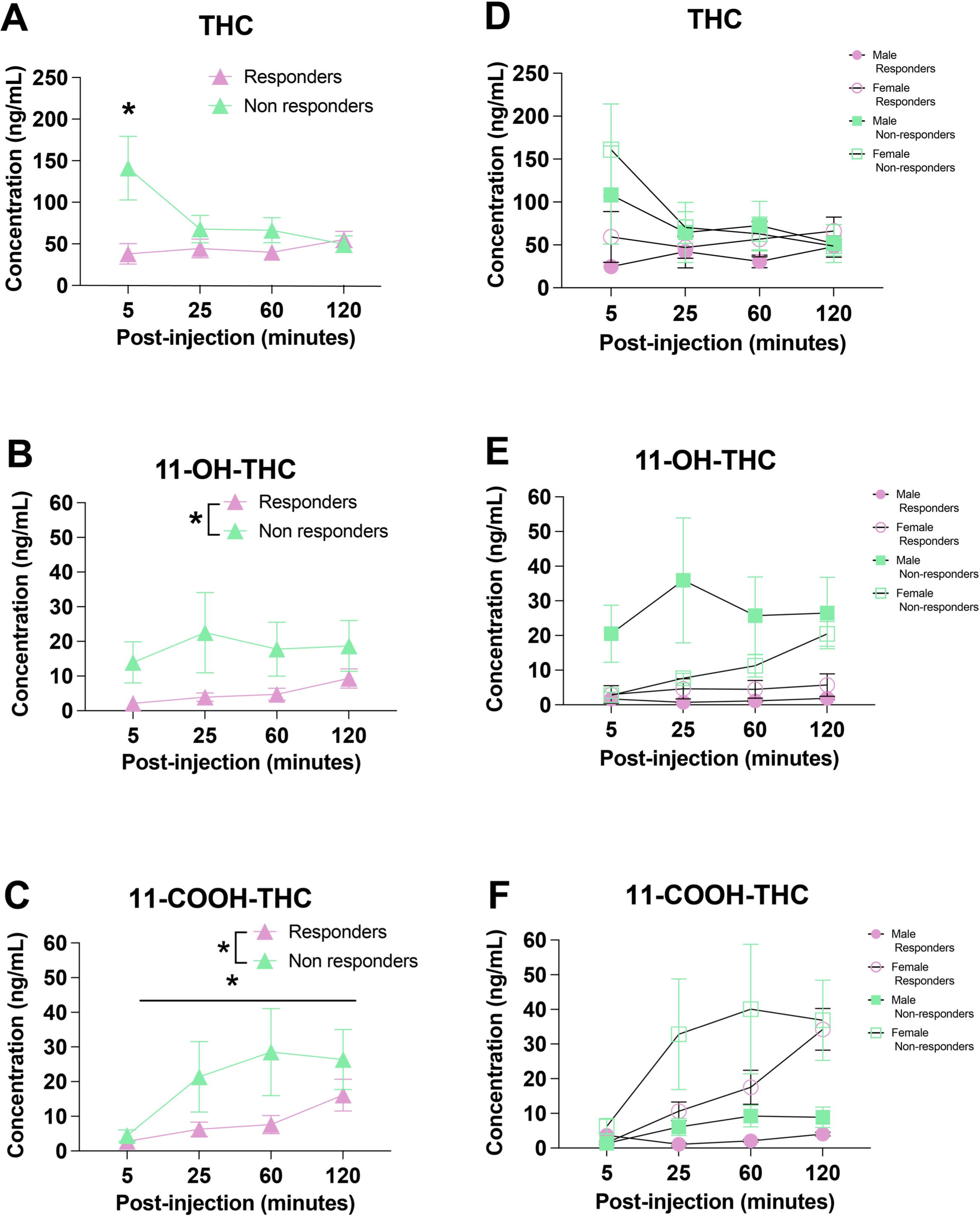
THC and its major metabolites plasma concentration in responders and non-responders following 3mg/kg THC administration. A) THC, B) 11-OH-THC, and C) 11-COOH-THC plasma concentrations over time collapsed by sex. D) THC, E) 11-OH-THC, and F) 11-COOH-THC plasma concentrations over time stratified by sex. Male responders n = 8, female responders n = 6, male non-responders n = 3, and female non-responders n = 5. All data are presented as mean (± SEM), * p < .05

**Figure 4.**
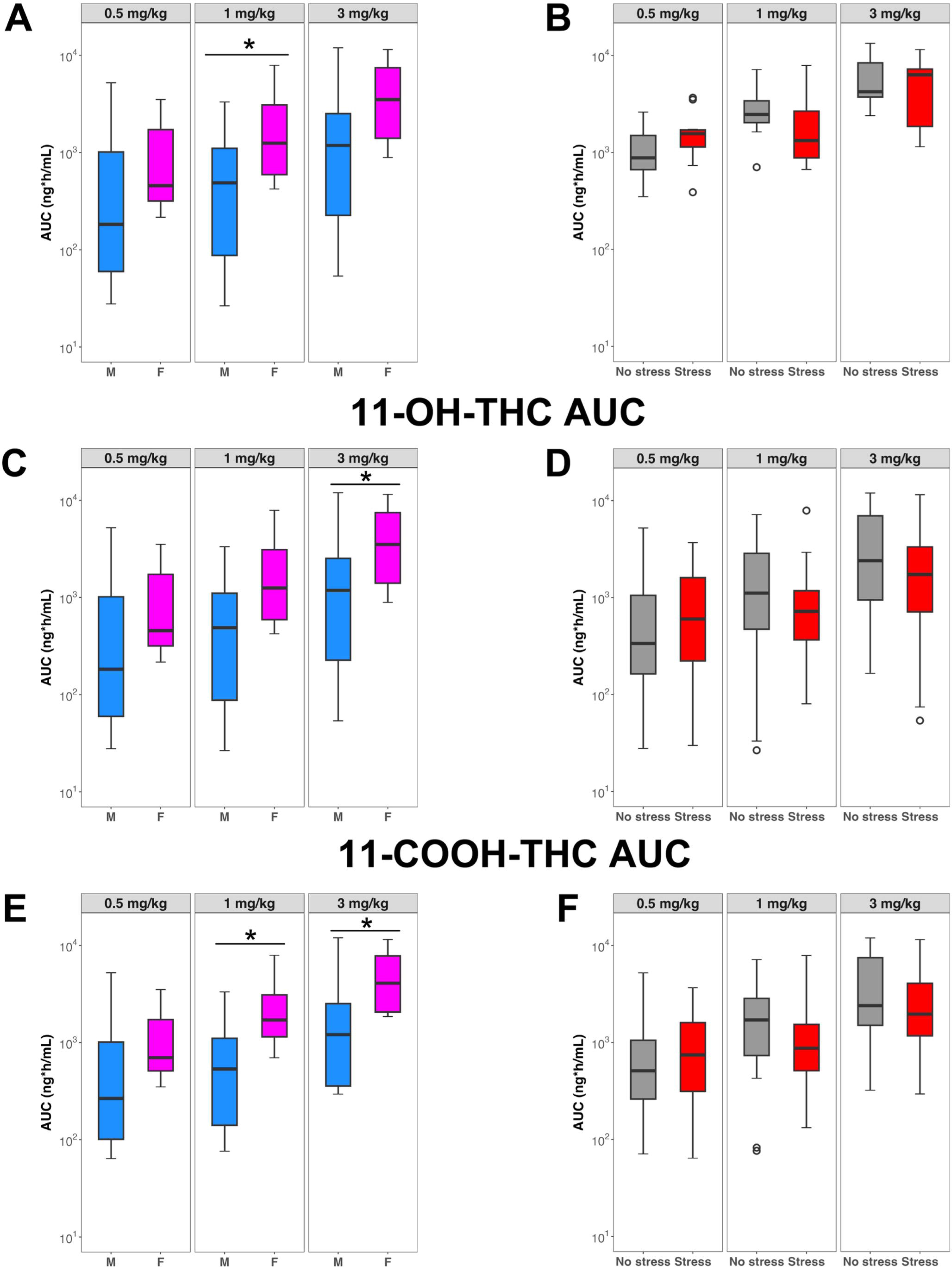
Distributions of THC and its major metabolites AUC values across each dose. A) THC, C) 11-OH-THC, and E) 11-COOH-THC AUC values stratified by sex across each dosing group. B) THC, D) 11-OH-THC, and F) 11-COOH-THC AUC values stratified by stress group across each dosing group. Boxplots are medians with 25th and 75th percentiles. Whiskers display the data points within 1.5-times the interquartile range below and above the lower and upper quartile values, respectively. * p < 0.05 via Wilcoxon rank-sum test. p < 0.05 via Kruskal-Wallis and Dunn’s test with Bonferroni correction.

However, stratifying by sex resulted in a low number of animals within each group, preventing our ability to statistically compare male and female responders and non-responders. Therefore, we combined males and females to analyze responders and non-responders. In doing so, a 2-way ANOVA revealed a main effect of time [F (3, 57) = 4.879, *p*=0.0043], a main effect of group [F (1, 21) = 5.303, *p* = 0.0316] as well as a time x group interaction [F (3, 57) = 7.707, *p* = 0.0002]. Interestingly, non-responders showed elevated THC plasma concentrations 5 minutes post 3mg/kg THC administration compared to responders (Figure 3B). From here, plasma concentrations of THC were similar between responders and non-responders at the rest of the timepoints collected. Next, we investigated the concentration of the major psychoactive THC metabolite, 11-OH-THC, and found a main effect of group [F (1,14) = 7.336, p = 0.0170] with the non-responders expressing significantly greater levels of the metabolite (Figure 3D). There were no effects of time [F (3, 42) = 1.666, *p =* 0.1889] or a time x group interaction [F (3,7) = 1.643, *p* = 0.2646] detected, indicating that non-responders had elevated 11-OH-THC across timepoints measured. Analysis of the non-psychoactive metabolite, 11-COOH-THC in our responders and non-responders revealed a main effect of both group [F (1,14) = 6.028, *p* = 0.0278] and time [F (3, 42) = 6.331, *p* = 0.0012], but no interaction [F (3,5) = 3.048, *p* = 0.1307] (Figure 3F), indicating that non-responders also showed elevated plasma 11-COOH-THC concentration compared to responders, and that levels in both groups increased with time. The elevated plasma concentration suggests that the non-responders metabolize THC at a faster rate than the responders. Overall, these data suggest that the lack of responding on the working memory task may be related to higher plasma THC and metabolite concentrations observed in the non-responders.

## 4. Discussion

The current study explored the relationship between plasma concentrations of THC and its major active and inactive metabolites with cognitive behavioral endpoints. Additionally, we explored whether stress modulated the pharmacokinetic profiling and cognitive behavior. We demonstrated that there was no alteration in working memory performance following acute restraint stress exposure. These results were not in line with our initial hypothesis, which predicted that stress would impair working memory performance. Previous research in rodent models has shown that exposure to acute stress can impair cognitive functioning in prefrontal cortex-dependent tasks, including the delayed-match-to-sample working memory test ^25,26^. On the contrary, non-prefrontal cortex-dependent memory tasks, such as tasks that are hippocampus or amygdala focused, have shown to be resilient to acute stress effects^25,27^. It is not clear why no effects of stress were observed here, but it is possible that our female population could be more resilient to the effects of stress as has been reported in other studies^28–30^.

Another possibility is that the timing of the stress exposure 30 minutes prior to working memory testing allowed for sufficient recovery of the physiological stress response. This interpretation is in support of previous studies showing that the effect of stress on prefrontal cortex-dependent tasks may lead to impairment soon after stress exposure and may return to baseline performance after time, due to the effect being driven in part by peak catecholaminergic and corticosteroid activity, which weakens normal PFC functioning but returns to more homeostatic levels post-acute stress ^31–33^. Additional studies are needed to fully understand the effect of stressor timing on cognitive functioning.

When administered an acute dose of either 0.5,1, or 3mg/kg THC 30 minutes prior to working memory testing, there was no effect on working memory performance in the rats that completed a sufficient number of trials to make accurate performance estimates. Our results are contradictory from results found in a study by Blaes et al., (2019) where they found that acute administration of 3mg/kg THC decreased working memory performance accuracy in both male and female rats. Although we did not see an effect of stress and/or THC on working memory performance, what we did find is that there were individual differences in task engagement that emerged following acute THC administration, specifically at the high dose of THC (3mg/kg). This data suggests that THC did impact performance on the task, but it was through effects on locomotion and/or motivation, not necessarily cognitive functioning. The lack of responsiveness during the task may be attributable to the recent work exploring cannabis and the amotivation hypothesis, which explores the relationship between cannabis use and lack of motivation^2,34,35^. It is also possible that prior studies reporting cognitive deficits after THC administration were affected by performance deficits in some individuals. These adverse effects on performance align with previous literature showing that higher concentration THC products can lead to greater adverse consequences following use^36–38^.

When exploring what may be mediating these individual differences in the willingness to respond, we found that there were significant differences in the pharmacokinetic and metabolic properties of THC in the responders vs non-responders. The differential profiling of these two groups suggests that plasma THC concentrations may be utilized to predict individual differences in sensitivity to the effects of THC. Specifically, high plasma concentrations of THC and its active metabolite 11-OH-THC may be related to the impact of THC on behavior. Individual differences in the pharmacokinetic and metabolic effects of THC have largely been overlooked when assessing neurocognitive or intoxicating effects of THC^16,22^. Our results are amongst the first to showcase the importance of taking into account these individual differences when assessing behavior. Because THC demonstrates a biexponential concentration-time profile due to rapid tissue distribution followed by slower metabolism and elimination^24,40^, the AUC estimates reported herein will underestimate the true total exposure due to sample collection ceasing prior to achieving pseudoequilibrium. Despite these limitations, our pharmacokinetic results align with previously published findings. Specifically, our results show at least 2-fold higher female median AUC estimates for both metabolites in each dosing group. Although only 11-OH-THC in the 3 mg/kg dosing group, and 11-COOH-THC in both the 1 and 3 mg/kg dosing groups showed statistical significance, an increased sample size may have revealed statistical significance all dosing groups. Showing similar trends, Torrens et al. found the AUC estimates for both 11-OH-THC and 11-COOH-THC to be significantly higher in females compared to males after acute 5 mg/kg i.p injections of THC^41^. There is disagreement in the literature regarding sex differences in THC exposure after i.p. injection. Baglot et al. found consistently higher THC concentrations in male rats compared to those in female rats up to 240 minutes after a 2.5 mg/kg acute i.p injection^24^. Additionally, our lab has found that following intravenous THC self-administration of equivalent doses, males exhibit greater plasma THC concentration compared to females. Conversely, Torrens et al. found no difference in parent drug exposure between sexes up to 480 minutes after a 0.5 or 5 mg/kg acute i.p injection^41^. Our findings in the 0.5 mg/kg and 3 mg/kg groups agree with the latter, but we noted that female THC AUC estimates were significantly higher than males in the 1 mg/kg dosing group.

## 5. Conclusion

In conclusion, our data showcases that higher plasma THC and metabolite concentrations may be associated with levels of adverse outcomes. Our data highlight the importance of studying individual differences in the effects of cannabis use and suggest that plasma THC concentrations may be predictors of the behavioral effects of cannabis. Understanding factors that may predict an individual’s susceptibility to the beneficial or adverse effects of cannabis will aide in effectively making individualized decisions on the use of cannabis medicinally. Additionally, with the push for the legalization of cannabis products, discovering factors that mediate one’s resiliency or vulnerability to the negative effects of THC is of increasing importance. Ongoing studies are needed to expand upon the current findings by investigating the relationship between stress, cannabinoid 1 receptor binding, and working memory performance.

## Supplementary Materials

Includes figures S1-S4

## Acknowledgements

Phoenix WinNonlin was generously provided to the University of Pittsburgh School of Pharmacy by Certara, Inc., through the Center of Excellence Program for academic institutions.

## CRediT authorship contribution statement

**Teneisha Myers:** Investigation, Formal analysis, Data curation, Visualization, Validation, Writing – original draft, Writing – review and editing **Noah Neverette:** Formal analysis, Data curation, Visualization, Validation, Writing – original draft, Writing – review and editing **Aaron Devanathan:** Conceptualization, Methodology, Resources, Writing – review and editing, Supervision, Project administration, Funding Aquisition **Mary Torregrossa:** Conceptualization, Methodology, Resources, Writing – review and editing, Supervision, Project administration, Funding Aquisition

