## Supplemental Figures for "Individual differences in the pharmacokinetic profiling of Δ9-THC may be associated with differential motivational effects of Δ9-THC on cognitive performance"

**Supplemental Figure 1.** Concentration-time curves in males (blue line) and females (magenta line). Closed circles represent medians and whiskers represent 25th and 75th percentiles. The black dashed line represents the lower limit of quantification for all analytes (0.1 ng/mL).

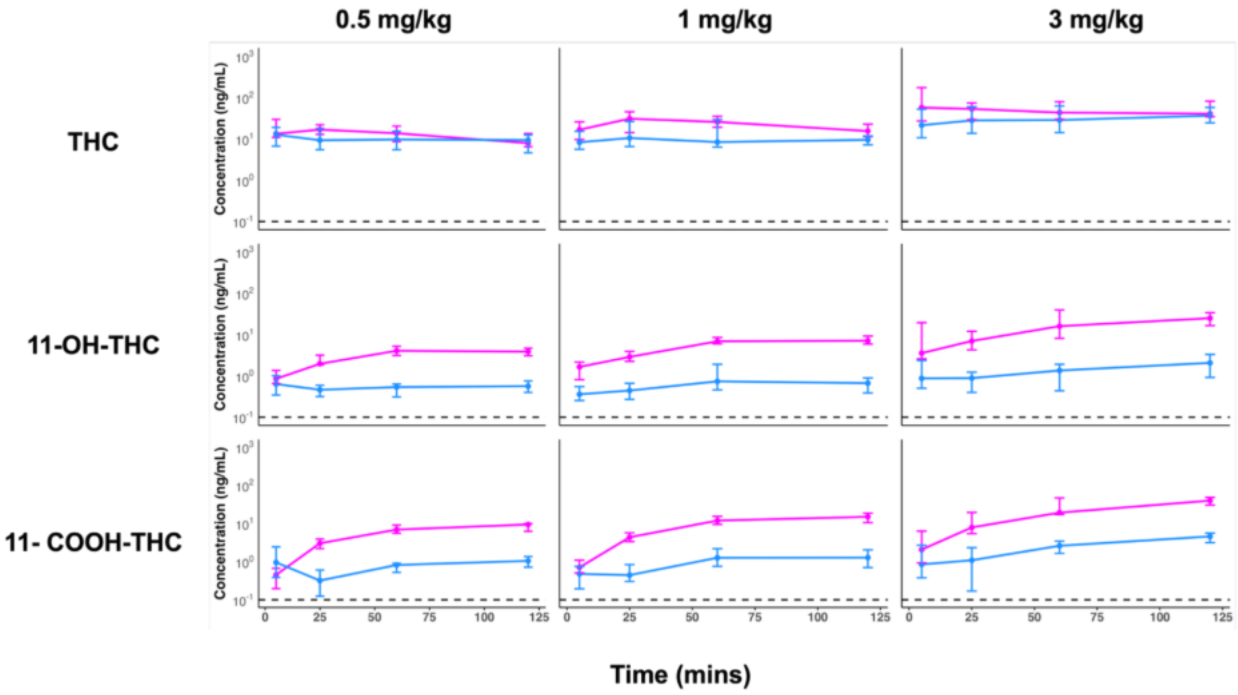

**Supplemental Figure 2.** Concentration-time curves in stressed (red line) and non-stressed controls (black solid line). Closed circles represent medians and whiskers represent 25th and 75th percentiles. The black dashed line represents the lower limit of quantification for all analytes (0.1 ng/mL).

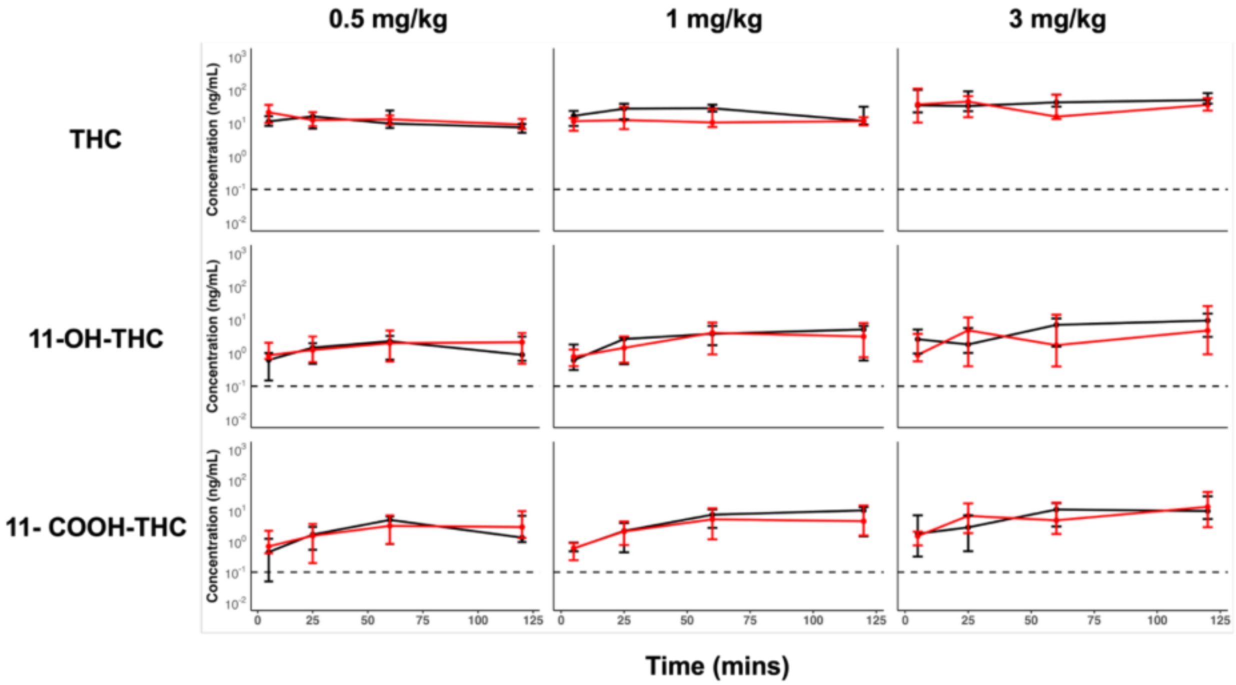

**Supplemental Figure 3.** Concentration-time curves in stressed males (blue line), non-stressed males (green line), stressed females (maroon line), and non-stressed females (magenta line). Closed circles represent medians and whiskers represent 25th and 75th percentiles. The black dashed line represents the lower limit of quantification for all analytes (0.1 ng/mL).

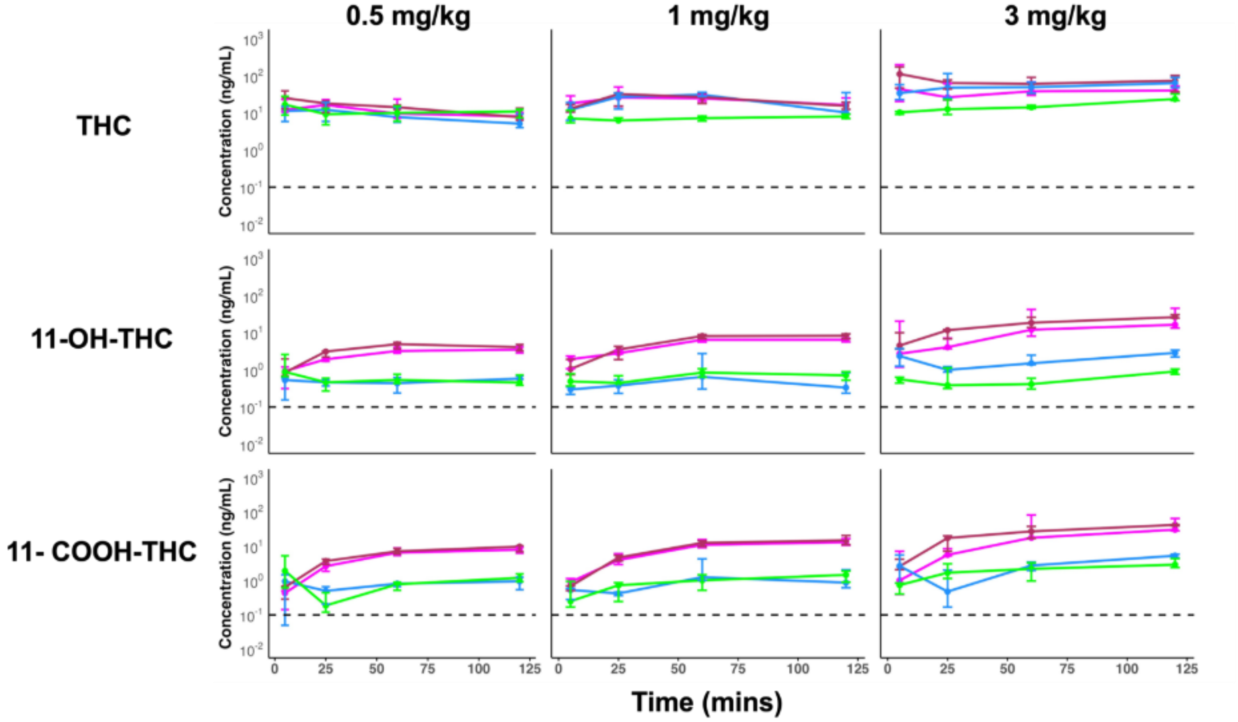

**Supplemental Figure 4. Distributions of THC and its major metabolites AUC values between stressed males (green), stressed females (maroon), non-stressed males (blue), and non-stressed females (magenta) across different dosing groups.** Boxplots are medians with 25th and 75th percentiles. Whiskers display the data points within 1.5-times the interquartile range below and above the lower and upper quartile values, respectively. * $p < 0.05$ via Kruskal-Wallis and Dunn's test with Bonferroni correction.

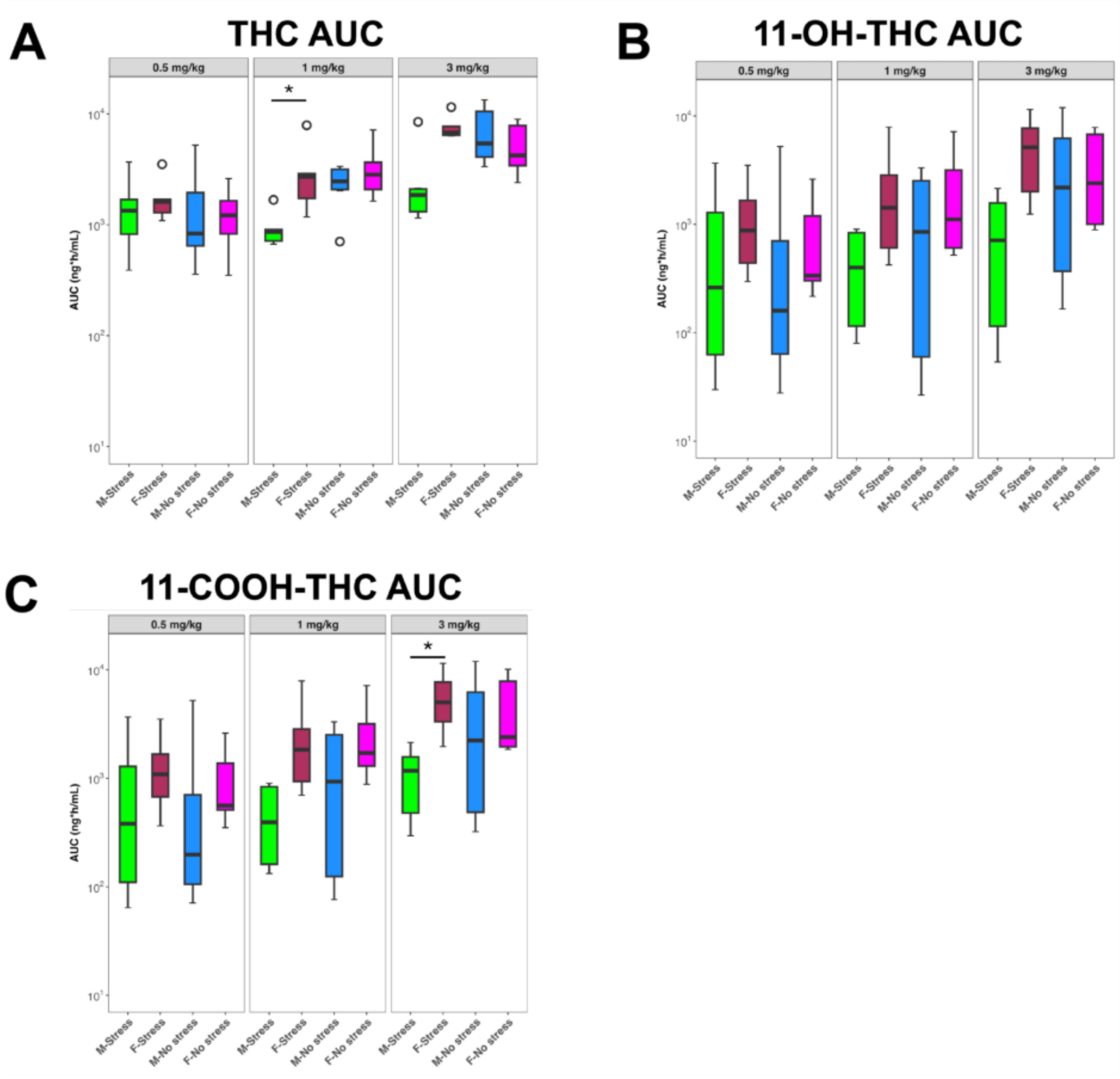
